# Ancient genomes reveal early Andean farmers selected common beans while preserving diversity

**DOI:** 10.1101/791806

**Authors:** Trucchi Emiliano, Benazzo Andrea, Lari Martina, Iob Alice, Vai Stefania, Nanni Laura, Bellucci Elisa, Bitocchi Elena, Xu Chunming, Jackson A Scott, Lema Verónica, Babot Pilar, Oliszewski Nurit, Gil Adolfo, Neme Gustavo, Michieli Catalina Teresa, De Lorenzi Monica, Calcagnile Lucio, Caramelli David, Star Bastiaan, de Boer Hugo, Boessenkool Sanne, Papa Roberto, Bertorelle Giorgio

**Affiliations:** Department of Life and Environmental Sciences, Marche Polytechnic University, Ancona, Italy; Department of Life Sciences and Biotechnology, University of Ferrara, Ferrara, Italy; Department of Biology, University of Firenze, Firenze, Italy; Department of Agricultural, Food, and Environmental Sciences, Marche Polytechnic University, Ancona, Italy; Center for Applied Genetic Technologies, University of Georgia, Athens, Georgia, USA; Universidad Nacional de Córdoba, Córdoba, Argentina; Consejo Nacional de Investigaciones Científicas y Técnicas, Argentina; Instituto de Arqueología y Museo, Universidad Nacional de Tucumán, Tucumán, Argentina; Instituto de Evolución, Ecología Histórica y Ambiente (CONICET & UTN FRSR), San Rafael, Argentina; Museo de Historia Natural de San Rafael, Argentina; Instituto de Investigaciones Arqueológicas y Museo “Prof. Mariano Gambier” - UNSJ, San Juan, Argentina; Museo Arqueológico de Cachi, Cachi, Argentina; Department of Mathematics and Physics “Ennio De Giorgi”, University of Salento, Italy; Centre for Ecological and Evolutionary Synthesis, Department of Biosciences, University of Oslo, Oslo, Norway; Natural History Museum, University of Oslo, Norway

**Keywords:** Ancient DNA, plant genomics, domestication, genomic erosion, selection scan, sustainable agriculture

## Abstract

All crops are the product of a domestication process that started less than 12,000 years ago from one or more wild populations [1, 2]. Farmers selected desirable phenotypic traits, such as improved energy accumulation, palatability of seeds or reduced natural shattering [3], while leading domesticated populations through several more or less gradual demographic contractions [2, 4]. As a consequence, erosion of wild genetic variation [5] is typical of modern cultivars making them highly susceptible to pathogens, pests and environmental change [6,7]. The loss of genetic diversity hampers further crop improvement programs to increase food production in a changing world, posing serious threats to food security [8,9]. Using both ancient and modern seeds, we analyzed the temporal dynamic of genetic variation and selection during the domestication process of the common bean (*Phaseolus vulgaris*) that occurred in the Southern Andes. Here we show that most domestic traits were selected for prior to 2,500 years ago, with no or only minor loss of whole-genome variation. In fact, i) all ancient domestic genomes dated between 600 and 2,500 years ago are highly variable - at least as variable as a modern genome from the wild; the genetic erosion that we observe in modern cultivars is therefore a recent process that occurred in the last centuries; ii) the majority of changes at coding genes that differentiate wild and domestic genomes are already present in the ancient genomes analyzed here. Considering that most desirable phenotypic traits are likely controlled by multiple polymorphic genes [10], a likely explanation of this decoupling of selection and genomic erosion is that early farmers applied a relatively weak selection pressure [2] by using many phenotypically similar but genomically diverse individuals as breeders. Selection strategies during the last few centuries were probably less sustainable and produced further improvements focusing on few plants carrying the traits of interest, at the cost of marked genetic erosion.

## Introduction

The onset of domestication by early farmers has been suggested as the period during which the most intense genetic bottleneck affecting genome-wide diversity occurred [4,11]. Yet, artificial selection, possibly at different rates in different times, was likely a continuous process that produced both landraces and, more recently under modern breeding programs, high-yielding and more resistant elite cultivars [2,12,13]. Understanding when the majority of genetic diversity was lost, and how such loss is related with the intensity of the selection process, is not only relevant from an evolutionary perspective but it will also help planning more sustainable breeding options to revert or mitigate crop’s genetic erosion [14,15]. We directly address these questions by analyzing both modern and ancient genomes of common bean from South America. Common bean constitutes one of the major sources of vegetable proteins worldwide. The wild progenitor of common bean originated in Mesoamerica and colonized South America along the Andes [16,17]. This natural colonization process, dated by whole genome analysis between 146,000 and 184,000 years ago [17], was accompanied by an intense and long bottleneck, as supported by the higher genetic diversity observed in the Mesoamerican as compared to the Andean gene pool [18]. Domestication of common bean occurred independently more or less simultaneously in both Mesoamerica and the Andes ca. 8,000 years ago, followed by divergence into distinct landraces due to drift, local selection, and adaptation [16].

## Results and Discussion

We obtained whole genome *shotgun* data from 30 ancient common bean seeds representing nine archaeological sites in north and central western Argentina (Supplementary Information S1-2,4, Supplementary Figure S1, Supplementary Table S1). All 30 seeds were dated between 2500 and 600 years before present (yrs BP) by AMS radiocarbon dating (Supplementary Information S3). Initial screening showed remarkable DNA preservation, with on average 44% (SD = 12%) endogenous DNA mapped to the common bean reference genome [17] (Supplementary Table S1). Short fragment lengths (average = 65, SD = 22 bp) were consistent with those expected for degraded DNA. The presence of deamination patterns at the end of the reads varied from 0% to 30% C to T and G to A misincorporations, with half of the seeds showing percentages below 10% (Supplementary Figure S2). Damage patterns were partially explained by archaeological site’s locations: all seeds from sites at >2,500 meters above sea level showed less than 10% base misincorporations (with many close to or at zero), whereas the majority of seeds from lower altitude exhibited levels above 15% (Supplementary Figure S3). The observed pattern suggests that both favorable environmental conditions and the structure of the seed (*e*.*g*., presence of an external cuticle) likely resulted in good DNA preservation. A subset of the specimens (Figure 1a,b) was selected for further sequencing based on DNA preservation and representation of age and locations, resulting in a final dataset of 15 ancient bean genomes sequenced from 4X to 18X coverage (Supplementary Table S1). Whole-genome data from modern common bean accessions (domesticated landraces/cultivars: 9 Andean, 3 Mesoamerican; wild: 1 Andean, 1 Mesoamerican) and a closely related species (*P. hintonii* 1 accession) were also included in the analyses (Supplementary Information S6; Supplementary Table S2).

**Figure 1.**
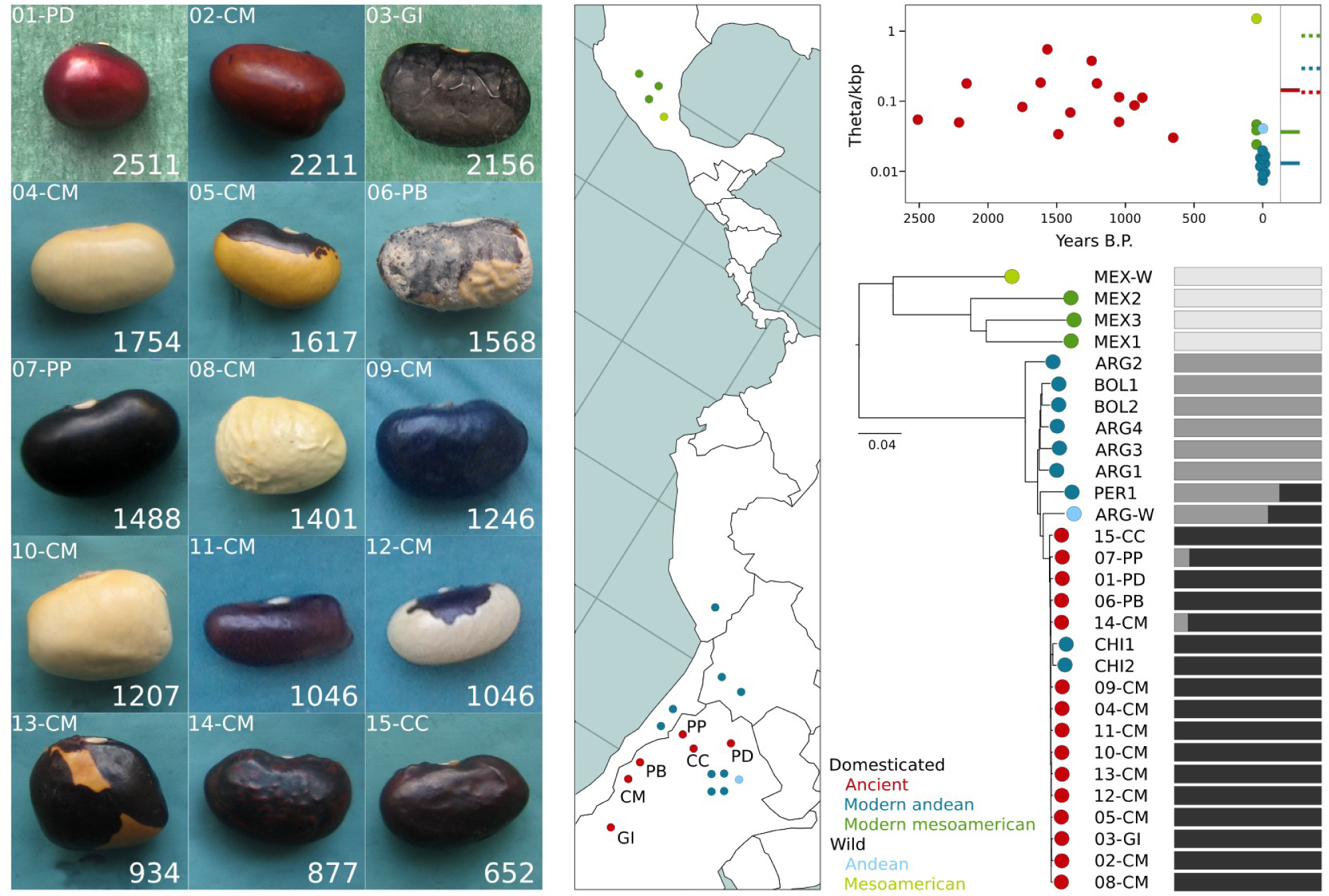
Diversity in ancient and modern common bean. **A)** Pictures of common bean seeds used for high-coverage whole-genome sequencing: sample ID with two letters referring to the archaeological site of origin as shown in panel B is reported in the top-left corner of the pictures (see Supplementary Information S1); AMS radiocarbon age in yrs BP is reported in the bottom-right corner. **B)** Map showing the locations of the archaeological sites where the seeds were retrieved (red) and the country of origin of the modern domesticated cultivars from South and Central America (blue and green, respectively) and the wild ones (pale blue, pale green). Modern seeds location is only indicative of the country of origin **C)** Whole-genome diversity estimated as θ kbp-1 in individual seeds (solid circles), average of individual estimates per group (solid line), and in groups of seeds (dashed line) using only transversions in callable regions (see Supplementary Information S5 for details); only domesticated cultivars are included in the group estimates. **D)** Neighbor-Joining tree based on genetic distances and Admixture analysis both based on transversions only (one SNPs every 50 kb was used in the Admixture analysis) in callable regions; note that modern seeds from Chile cluster together with the ancient seeds.

Heterozygosity is higher in ancient than in modern domesticated Andean seeds. In particular, individual heterozygosity (estimated as the variation parameter θ every thousand nucleotides; Supplementary Information S7) is higher in all ancient seeds than in each of nine modern landraces/cultivars from the same Andean gene pool (Figure 1c, Supplementary Table S3) and average heterozygosity is more than ten times higher in ancient than in modern seeds (0.144 vs. 0.013 θ/kbp, respectively, Mann-Whitney U, *P*<0.001). The possible effect of decreased variation due to the selfing procedure applied to modern accessions in seed banks, was tested by pooling pairs of modern seeds and estimating θ/kbp per pair (Supplementary Information S8). As expected, variation in modern pairs increased compared to single seeds, but it is on average still three times smaller than that observed in ancient seeds (Mann-Whitney U test, *P*<0.005, Supplementary Figure S4; Supplementary Table S4). Considering that θ/kbp can increase in modern seed pairs also because our paired seeds belong to different landraces (i.e., to genetic structure among landraces and not to seed bank practice), we conclude that modern seeds are several magnitudes less variable than ancient seeds.

The only modern genome available from a wild Andean seed from Argentina [19] has a much lower genomic diversity than the wild Mesoamerican seed in our dataset (Figure 1c), and similar to the less variable ancient seeds. Although single seeds clearly represent only a fraction of the global variation, especially if populations are genetically structured, this result is a) consistent with the bottleneck that occurred during the natural colonization of the Andes from Mesoamerica [18] and b) it appears compatible with a minor effect of initial Andean domestication on genetic diversity. Moreover, the lack of a temporal trend across ancient seeds of different age (Figure 1c) indicates that genome-wide diversity was largely maintained by agricultural practices of Andean societies between 2500 and 600 years ago. A more consistent loss of diversity within each cultivar occurred more recently, certainly less than 600 years ago and likely in the last century [13]. Similarly, a very recent loss of diversity characterized the domestication history of horses [20]. The absence of data on seeds from the last 600 years hinders, however, a direct inference about the temporal dynamic in the last six centuries.

When estimating genetic variation in groups of modern Andean (0.294 θ/kbp) or Mesoamerican seeds (0.860 θ/kbp) we observe a substantial increase compared to average individual estimates (Figure 1c), indicating that modern genomic diversity is highly structured in different, highly homogeneous, cultivars. The abandonment and extinction of any modern cultivar would therefore result in the significant loss of its private fraction of the whole crop diversity [8,21]. On the contrary, the estimate of genetic diversity obtained by pooling all ancient individuals is very similar to the average of the 15 individual estimates (0.134 θ/kbp; Figure 1c), clearly indicating that ancient genomes belong to a single genomic pool (*i*.*e*., they are very similar both across time and archaeological sites) and that any handful of ancient seeds could recover a large fraction of the ancient crop diversity.

Three main genetic groups were revealed by different clustering methods (Supplementary Information S9) and a whole-genome NJ tree (Figure 1b; Supplementary Figures S5-S6). The largest divergence is observed between Mesoamerican and Andean genomes. The Andean seeds are then further partitioned into two less differentiated groups. All ancient seeds belong to the same genomic clade, indicating that the same or similar ancient landraces were in use in Argentina for about 2,000 years. Rather surprisingly, this clade also includes modern cultivars used today on the other side of the Andes, in Chile. The so-called Chilean race [22] therefore appears to be the direct descendants of the seeds used in Argentina before the Incaic conquest of the area occurred at the end of the 15^th^ century [23]. Modern Argentinian cultivars, instead, do not trace their ancestry back to the local ancient cultivar but were likely introduced in the area some time after 600 years ago.

A gene-by-gene selection scan (Figure 2a) was performed to discriminate early (>2,500 years ago) from late (<600 years ago) selection targets, by taking into account the different levels of genetic drift expected in the two time periods of different length (Supplementary Information S10). We found that: *i*) a selection signature is present in approximately 400 out of 27,000 genes tested (FDR<0.001); *ii*) selection affected many more genes (ca. 4.5 times more) in the early compared to the late phase (Fig. 2b), suggesting that recent improvement of cultivars strongly affected few genes, but the main genomic turnover occurred several thousand years ago; *iii*) genes selected in the early phase belong to functional clusters mainly related to glycerol metabolism, carbohydrate and sugar transport and metabolism, intracellular transport (endosome membrane), regulatory elements (nucleotide binding), modification of proteins, glycosylation, whereas recent selection mainly targeted traits involved in immunity and defense, regulatory elements and transmembrane transport (Figure 2c; Supplementary Table S5); *iv*) nine genes show a significant signature of both early and late selection and are mainly related to immune response. Evidence for selection on sugar biosynthesis genes at the early stages of domestication has also been found in ancient maize from 2000 and 750 years ago [24]. One of the functional clusters identified in the early selected genes includes the MYB-domain genes that are important regulatory elements of development, metabolism and responses to biotic and abiotic stresses [25], and have been suggested as causative of different phenotypic changes associated to shattering trait in common bean [26], maize and rice [4,27].

**Figure 2.**
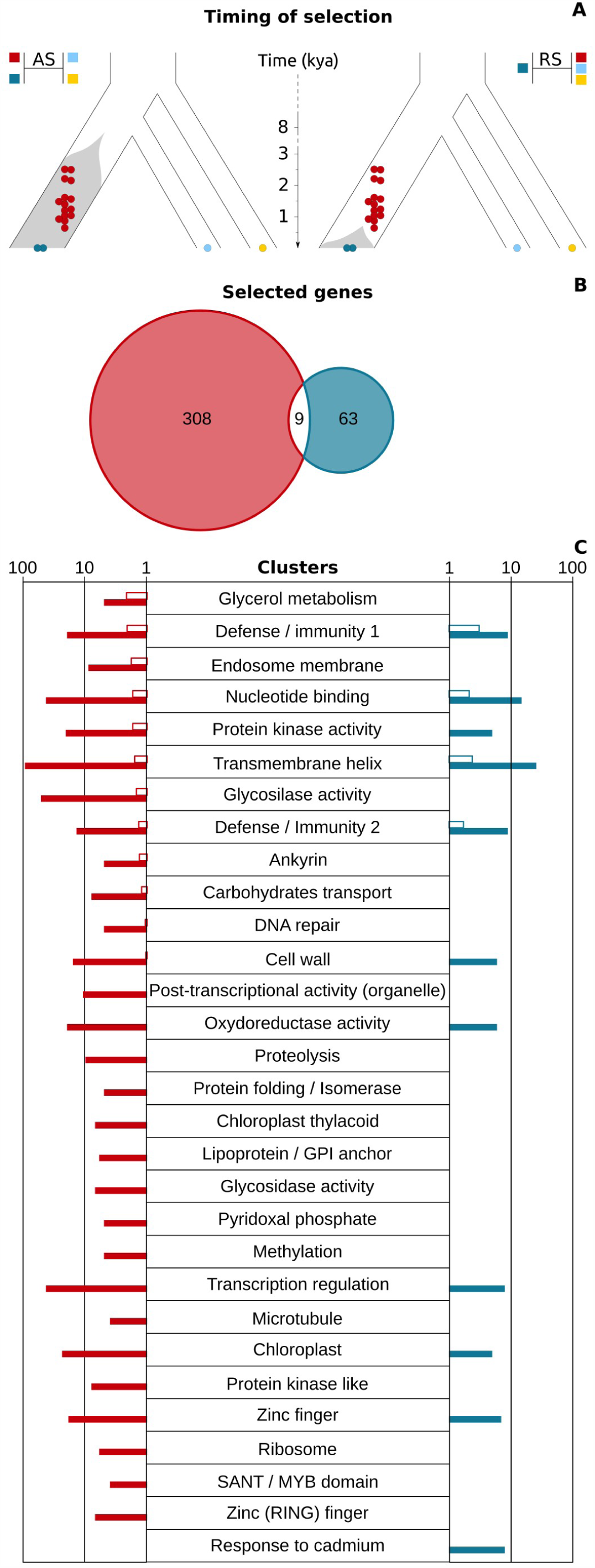
Timing of selection in common bean from South America. All genes (27,000) in the common bean genome were tested for a significant enrichment of fixed alternative alleles according to the topologies described below but taking into account different level of drift occurring during period of time of different length (see Supplementary Information S10 for details). **A)** Alternative topologies for a SNP: on the right (AS: ancient selection) the SNP is fixed for an alternative variant in both the ancient seeds from Argentina (red circles) and the modern cultivars from Chile (blue circles) as compared with the wild andean *P. vulgaris* seed (pale blue circle) and the outgroup (*Phaseolus hintonii*, yellow circle); on the left (RS: recent selection) the SNP is fixed for an alternative variant in the modern cultivars from Chile only as compared with the ancient seeds, the wild Andean and the outgroup; the gray shade represents the temporal fixation of an alternative allele; ancient seeds age as well as the putative divergence time between Andean domesticated and wild *P. vulgaris* is reported. Using the genome of the wild mesoamerican *P. vulgaris* as outgroup instead of *P. hintonii* produced largely similar results (see Supplementary Table S5). **B)** Number of genes significantly enriched with SNPs in RS (blue) and/or AS (red) topology (FDR<0.001, top 1% for the proportion of SNPs within gene ± 1kb, and with an *Arabidopsis thaliana* ortholog). **C)** Functional gene clusters (see Supplementary Information S10) in RS (blue) or AS (red) enriched groups (the complete list of genes can be found in Supplementary Table S5). Filled bars: max gene count in the cluster; empty bars: enrichment score of the cluster (only scores >1 are reported).

## Concluding remarks

Genomic studies of ancient DNA are much less numerous in plants compared to animals remains [28-29], but have already provided fundamental insights into the process of domestication [24,30-34]. The dynamic of this process is still debated and likely not the same in all species, but, as recently summarized [2], the consensus is that domestication was slow and gradual, and selected traits emerged at the cost of a decline in population size and genetic variation. Here we contribute to this debate with the first study on ancient bean genomics, and we suggest an alternative scenario under which selection and loss of variation are decoupled in this species. First of all, our sequencing effort shows that common bean seeds, and likely legume seeds in general, can be an excellent source of high quality ancient DNA, especially if seeds are preserved in favorable climatic conditions (*i*.*e*., cold and dry environments). Second, our data reveals that in common bean early agriculture was more sustainable for genomic diversity compared to more recent breeding. At the same time, however, assuming that the genetic changes we infer correspond at least in part to phenotypic changes, early agriculture was also very efficient in selecting most of the desirable traits now typical of this crop. We hypothesize that these patterns are the result of the larger number of seeds, likely all displaying the selected trait(s) but heterogeneous in the rest of the genome, used by early farmers as founders in each generation. Initial improvement of the common bean was therefore based on serial soft sweeps, as has also been suggested for the domestication of maize [10]. Encompassing several thousand years, and likely assisted by cultivar exchanges and hybridization with wild plants [2,22,35-36], such breeding practice allowed many traits to be selected for without the significant loss of genomic variation that has occurred in more recent times.

## Acknowledgments

This study was supported by CUIA (Consorzio Universitario Italiano per l’Argentina, V^ Research Program), the Bean_Adapt (Era-Caps) project, the internal grants from the University of Ferrara, the Marche Polytechnic University, and the University of Firenze, the Italian Ministry of Education, University and Research (project “Dipartimenti di Eccellenza 2018-2022”), the Swedish Research Council grant VR-UF E0347601 and the Norwegian Research Council grant 262777. We also thank CONICET (Consejo Nacional de Investigaciones Cientificas y Técnicas) and the Institutions and Museums in Argentina for their support in the fieldwork, for recovering the archaeo-botanical specimens and for their investigation of archaeological sites.

## Data accessibility

Sequenced raw reads from ancient samples been made publicly available as NCBI Bioproject (ID: PRJNA574560). Unpublished modern sample genomic data are available upon request from ERA-CAPS funded project “BeanAdapt: The Genomics of Adaptation during Crop Expansion in Bean”, coordinated by Roberto Papa, Marche Polytechnic University.

## SUPPLEMENTARY METHODS AND RESULTS

### S1. Ancient beans collections and archaeological context

Ancient bean seeds were selected from five museum collections in Argentina: Museo de La Plata (La Plata, Buenos Aires, Argentina); Museo de Historia Natural de San Rafael (Parque Mariano Moreno, San Rafael, Argentina); Instituto de Investigaciones y Museo Arqueológico “Prof. Mariano Gambier”, Universidad Nacional de San Juan (San Juan, Argentina); Instituto de Arqueología y Museo, Universidad Nacional de Tucumán (Tucumán, Argentina); Museo Arqueológico Pio Pablo Díaz (Cachi, Salta, Argentina).

Seeds were originally collected from nine archaeological sites located in different geographical regions of north and central-western Argentina (Figure 1a, main text; latitude and longitude are available on request). A brief description of each site is reported below. Codes for each seed, referring to Figure 1b (main text) and Figure S1, are reported in Table S1.

#### Los Morrillos, Río Salado, Cerro Calvario and Punta del Barro (San Juan sites)

The San Juan sites are in the two west Departments of the Province of San Juan, Iglesia (to the north) and Calingasta (to the south). They are in a southern Andean area that extends between 28º25’S and 32º33’S on the arid eastern slope of the Andes. This territory includes (from west to east) the “Cordillera de los Andes”, two wide valleys between 1,900 and 1,600 m asl and another system of high altitude but of older age than the mountain range called “Precordillera of La Rioja, San Juan and Mendoza”; its maximum summits constitute the eastern limit of these departments. The first evidences of agricultural and breeding (*Lama glama*) activity did not originate in the area, but were introduced together with the domesticated species. The human groups that introduced and developed agriculture were located in favorable small sites corresponding to the exit of the cordilleran streams in the great plain of the high foothills (approximately 2,900 m asl). Such human groups cultivated in summer, gathered wild fruits and ñandu eggs, and entered the high Andean valleys for the hunt of the guanaco (*Lama guanicoe*). They mostly inhabited natural caves in the area. Examples are Los Morrillos (caves 2 and 3) and Río Salado. In the case of Los Morrillos caves, the thick level with agriculture overlapped with an older one of hunter gatherers. However, in the case of the Salado River, the cave only had a thick level with agriculture. The same groups, with some foreign influences, later descended towards the valleys at lower altitude and settled in open areas with the possibility of irrigation from small springs or streams. This is the case of the Cerro Calvario sites in the Calingasta Valley and Punta del Barro in the north of the Iglesia Valley (Basurero Norte, Basurero 3). Finally, the groups developed a large-scale agricultural system based on irrigation by means of large channels connected to the rivers, as is the case of the Punta del Barro “1º canal” (Gambier 1977, 1988, 2000, Michieli 2015, 2016, Roig 1977).

#### Pampa Grande

Pampa Grande is an archaeological locality including seven caves explored in the 1970s. It is located in the Las Pirguas mountain range (Salta province), between 2,500 and 3,000 m asl in a montane grassland and forest landscape corresponding to the highest zone of the *Yunga* biogeographical province. Ancient remains are mainly related to funerary contexts and few occupational areas, corresponding to Candelaria groups. The age of these occupations range between 400 – 3,000 yrs BP and corresponds to the first agro-pastoralist societies in the area. Remains related with subsistence include domestic, wild and hybrid plant species of *Cucurbita maxima* and *Phaseolus vulgaris. Phaseolus lunatus, Lagenaria siceraria, Zea mays, Arachis hypogaea, Prosopis* spp., *Geoffroeae decorticans*, tubers, *Capsicum* spp. and other wild and weed forms were recovered. Remains of wild and domestic camelids were also recovered together with fish, rodent and bird bones (Baldini et al 2003, Lema 2010).

#### Puente del Diablo

Puente del Diablo (SSalLap20) archaeological site is a cave situated in the northern sector of the Calchaqui Valleys in a pre puna landscape (Salta province). It was excavated in the 1970s and includes both domestic and funerary contexts. This is a multicomponent site with burials dating back to 10,000 yrs BP, remains corresponding to burials and temporal occupations of hunter gatherers of ca. 3,000 yrs BP and also from ca. 2,000 yrs BP corresponding to the first agropastoralist societies in the area. From these last occupations, remains of *Cucurbita maxima, Prosopis* spp. and cactaceae were recovered together with remains of rodents (*Lagidium* sp.), *Cervidae* sp. and camelids (*Lama* sp.) (Lema, 2009, 2015).

#### Gruta del Indio

Gruta del Indio is an archaeological site located in Central Western Argentina, close to San Rafael city, in Monte fitogeographic province, at 700 m asl. Gruta del Indio was occupied by hunter-gatherers since the Pleistocene-Holocene transition ca. 10,500 years ago (Semper and Lagiglia 1968, Long et al 1999). These hunter-gatherer occupations were in the cave until at least ca. 1,900 yrs BP, when a strong domestic plant record appears in the archaeological record. This domestic archaeobotanical record include *Zea mays, Chenopodium quinoa, Cucurbita*, and *Phaseolus vulgaris* seeds, and *Zea mays* remains, as canes and starches (Semper and Lagilia 1968, Lagiglia 1999). This domestic plant context (the oldest in the region) is isolated in time and space, and is associated to human bones remains, indicating the use of the cave as a cemetery. This archaeological context was called “Atuel II culture” (Semper and Lagiglia 1968) and all radiocarbon dates are between 2,100 and 1,900 yrs BP (Gil et al 2014). The stable isotopes on human bone suggest for this context that the *Zea mays* was probably part of the diet but not as major component (Gil et al 2010). The *Phaseolus vulgaris* beans were found in a grass container holding ca. 500 grs. of seeds. Another grass container was full of *Chenopodium quinoa* seeds in the same archaeological mortuary context. After Atuel II, there is few evidence of human occupation and temporally close to arrive of Spanish (ca. 500 yrs BP).

#### Punta de la Peña

Punta de la Peña 9 is a residential site with stone-walled structures, blocky shelters, bedrock mortars, caches, open air activity areas, and rock art that was continuously occupied by agro-pastoralist farmers between ca. 2,000 to 400 yrs BP -post-hispanic time (Babot et al 2006, López Campeny et al 2017, Somonte and Cohen 2006). It comprises processing, discard, workshop, funerary and ritual areas spatially segregated. The site is located on the southern margin of the Las Pitas River at an elevation of 3,665 m asl in the Salty Puna desert (Catamarca province). Subsistence was based on llama (*Lama glama*) herding, vicuña (*Vicugna vicugna*) hunting, small-scale cultivation, and plant gathering and exchange. Wild, domesticated and weed plant macro and micro-remains belong to a number of native and foreign taxa as *Chenopodium quinoa, Solanum tuberosum, Oxalis tuberosa, Phaseolus vulgaris, Prosopis* sp., *Geoffroea decorticans, Zea mays, Amaranthus caudatus/A. mantegazzianus, Lagenaria siceraria, Cucurbita* sp., *Canna edulis* and Cyperace and Cactaceae species (Babot 2009, Rodríguez 2013). Alero 1 is a blocky shelter rich in well preserved plant remains from the first millennium DC (1364 ± 20 cal. yrs BP), placed in the sector I of the Punta de la Peña 9 site (Babot et al 2013).

#### Cueva de los Corrales

Cueva de Los Corrales 1 is an archaeological site located in the Quebrada de Los Corrales (El Infiernillo, Tucumán) at 3,000 m asl in a dry shrub / grassland landscape, corresponding to the highest zone of the Yunga biogeographical province (Oliszewski et al 2008). It is a cave that was occupied at different times throughout 2,400 years (ca. 3,000-600 yrs BP), from transitional time between hunter-gatherer groups to agro-pastoralist groups until purely agro-pastoralist time (Oliszewski et al. 2018). Remains related to subsistence include domestic camelids and dasiphodids; wild plants: *Prosopis nigra, Geoffroeae decorticans, Celtis tala* and *Trichocereus* sp.; cultivated plants: *Phaseolus vulgaris, Zea mays* and *Chenopodium quinoa* and wild *Cucurbita maxima* (Oliszewski and Arreguez 2015, Oliszewski and Babot 2014, Lema 2017). The specimen of *Phaseolus vulgaris* here presented has a date of ca. 650 yrs BP corresponding to the last occupations of CC1.

### S2. Seeds selection

Thirty ancient bean seeds (see Figure S1) were selected for preliminary screening based on the site locations and archaeological context. All the seeds had a clear domestic phenotype. After visual inspection, we selected seeds with different morphological features (e.g. coat color and dimension) and good preservation (e.g., less visible cracks or damages in the outer coat).

**Figure S1.**
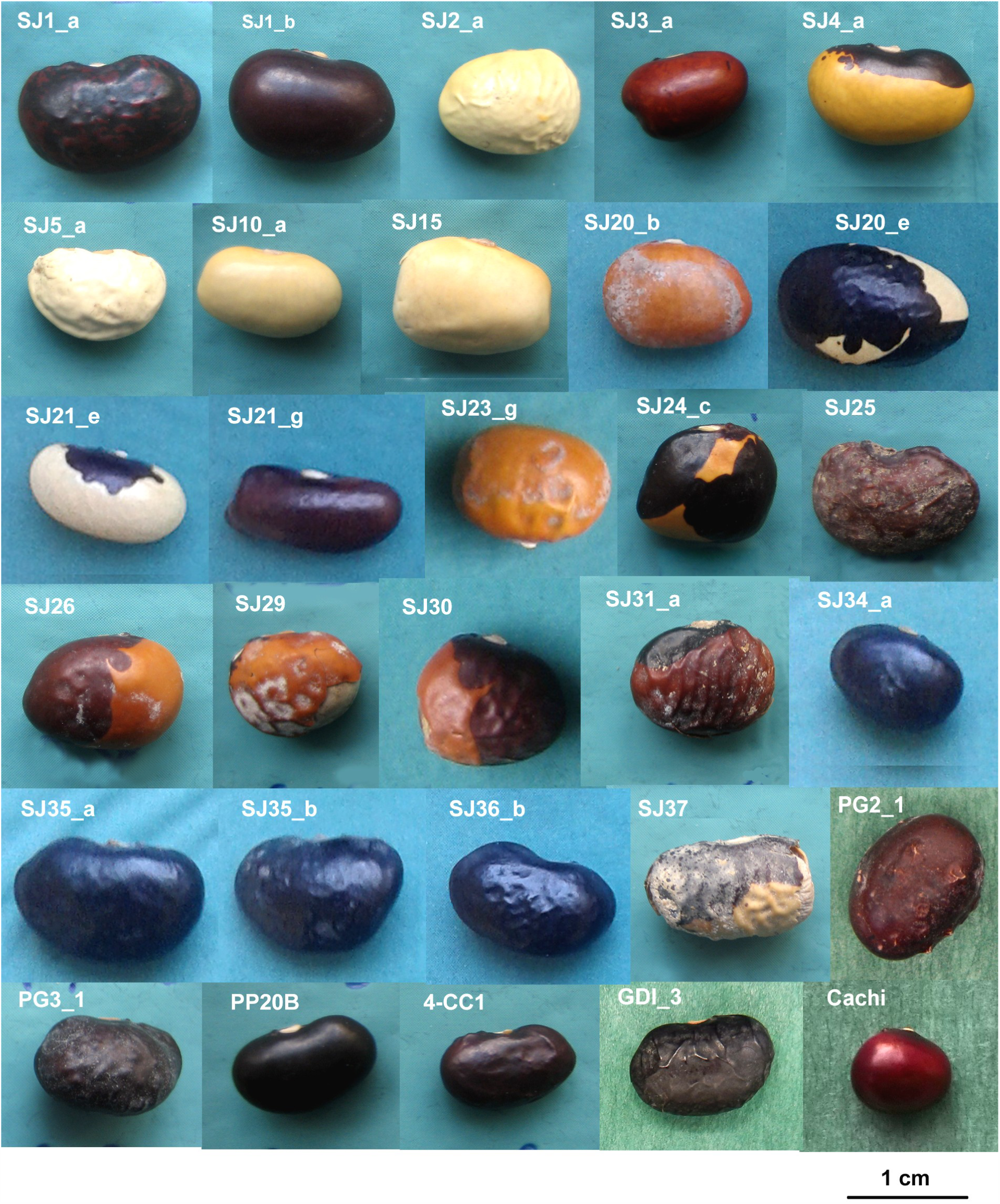
Pictures of the 30 seeds sequenced at low coverage for preliminary screening of aDNA content and preservation. Archaeological site information for each seed is reported in Supplementary Table S1.

### S3. AMS radiocarbon dating

A fragment of ca. ¼ of the cotyledon was removed from each seed and used for AMS radiocarbon dating at CEDAD, Center of Applied Physics, Dating and Diagnostics, Department of Mathematics and Physics “Ennio De Giorgi”, University of Salento. Conventional radiocarbon ages and calibrated ranges for 30 seeds are reported in Table S1. Calibration was performed using OxCal Ver. 3.10 based on atmospheric data (Reimer et al 2009). Distribution curves of the calibrated age for each seed, and additional details on the dating procedure are available on request.

### S4. Ancient DNA extraction and sequencing

Total DNA extraction and library preparation were performed at the Molecular Anthropology and Paleogenetic Laboratory of the Department of Biology, University of Florence, using facilities that are exclusively dedicated to processing ancient DNA and following all the necessary precautions to minimize contamination from exogenous DNA. One sample (09-CM) was extracted and a library built at the ancient DNA laboratory of the University of Oslo. Negative controls were included in each step of the molecular analysis. All seeds were photographed prior to sampling (Figure S1). After removing the cuticle, each seed was UV irradiated in a crosslinker for 45 minutes. The embryo and a portion of cotyledon were then pulverized by hand with mortar and pestle that had been decontaminated with bleach and UV. Half of the powder, was used for DNA extraction following a high-salt CTAB extraction protocol according to the Small Fragment Protocol of Qiagen DNeasy *mericon* Food Handbook. The amount of powder used in the DNA extraction ranged between 48 and 240 mg, depending on the size of the seed. Food Lysis Buffer was substituted with a homemade CTAB high salt extraction buffer according to the following recipe: 100mM Tris-HCl pH 8, 25mM EDTA pH 8, 4M NaCl, 2% CTAB, 0.3% β-mercaptoethanol. An aliquot of each extract was converted into a sequencing library with 7 bp P7 indexes according to Meyer & Kircher 2010. No UDG treatment was performed. The fill-in reaction was split in 4 aliquots that were amplified separately in 100µl reaction mixes with 0.25mM dNTPs mix (New England Biolabs), 0.30 mg/ml BSA (New England Biolabs), 2.5U Agilent Pfu Turbo Polymerase (Agilent Technologies) and 0.4µM of each primer. The following indexing PCR profile was performed by default on each library: 95°C 2’, [95°C 30’’, 58°c 30’’, 72°C 1’] x 14, 72°C 10’. DNA quantification and quality control were performed on a Agilent 2100 Bioanalyzer using the Agilent DNA 1000 kit, following manufacturer’s instructions. Libraries of extraction blanks all showed flat BioAnalyzer profiles, but were included in the pool of the first sequencing round (see below). The libraries were sequenced at the Norwegian Sequencing Centre, University of Oslo, Norway, where all samples were pooled in equimolar ratios and paired-end sequenced on the Illumina HiSeq 2500. First, samples were screened for DNA preservation by low-coverage sequencing. Based on the results of this first sequencing run and on sample characteristics (age, location, phenotype, mapping quality, age-related damage patterns), a subset of 18 samples was re-pooled and sequenced further on four Hiseq2500 lanes. Additionally, two samples (Cachi and SJ28) were sequenced on one lane each on the HiSeq 4000 platform. One sample (09-CM) was sequenced on the Hiseq 2500 in a different pool from all other beans. Based on the final results we further selected 15 samples with coverage higher than 4X for downstream population genomic analyses (Table S1).

### S5. DNA damage assessment, mapping and SNP calling

Paleomix v.1.2.12 (Schubert et al 2014), specifically designed for aDNA, was employed for the filtering and mapping of aDNA data. Adapter trimming, collapsing of overlapping mate-pairs, trimming of low quality bases and removal of reads shorter than 25 bp was performed with AdapterRemoval (Lindgreen et al 2012). Filtered reads were aligned to the *Phaseolus vulgaris* reference genome v2.1 (Schmutz et al 2014), downloaded from Phytozome (https://phytozome.jgi.doe.gov/pz/portal.html) using bwa aln (Li and Durbin 2009) and retaining reads with mapping quality >= 25. PCR duplicates were filtered using Paleomix rmdup_duplicates option and post-mortem damage was quantified in each sample (Figure S2) using MapDamage2.0 (Jónsson et al 2013). Indel-realigned bams were generated using the GATK Indel realigner (McKenna et al 2010). Realigned and filtered bam files were then indexed and sorted with Samtools-0.1.19 (Li et al 2009).

A descriptive evaluation of endogenous DNA preservation in the ancient bean seeds, as a function of the age of the sample and the site location is reported in Figure S3. Altitude, and not age, seems to be a good predictor of deamination pattern.

All following analyses including both ancient and modern (see section “Modern accessions genomic data”) samples were based on transversions only, since transitions are more affected by post-mortem damages than transversion and could reflect DNA damage rather than true DNA variation (e.g., Hofreiter et al 2012). The following regions were analyzed separately for the Admixture and NeighborJoining analyses (see below): *i*) *callable* regions, defined as the whole genome excluding repeated regions; *ii*) *neutral* regions defined as callable regions excluding genes (both exons and introns) and 10kb upstream and downstream each gene and *iii*) *exons*. The estimation of Watterson’s Theta (θ_w_) and the FineStructure analysis were performed on *callable* regions only (see below).

**Figure S2.**
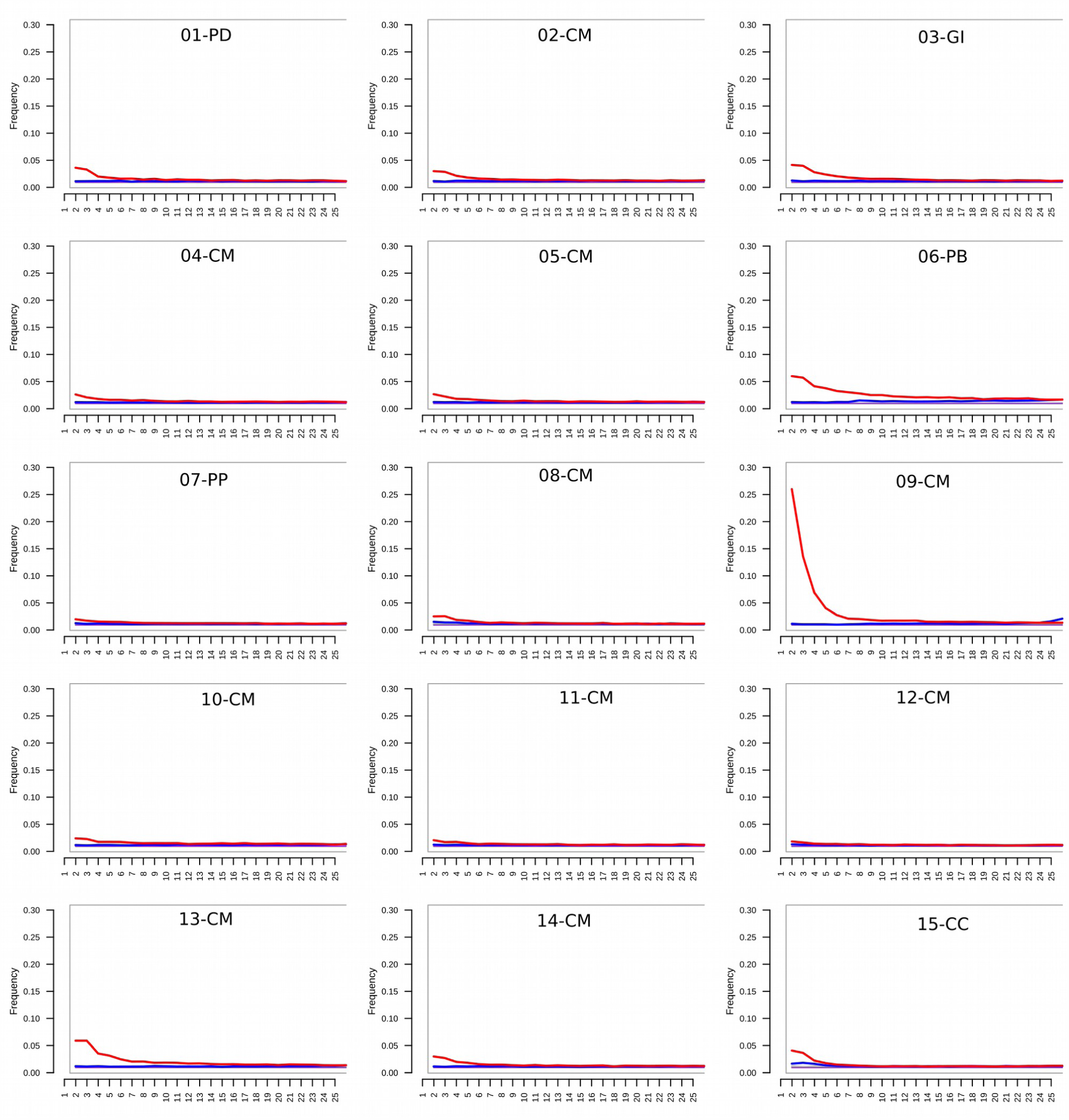
Ancient DNA damage plots for the 15 seeds included in the final analyses. Plots were generated using MapDamage 2.0.

**Figure S3.**
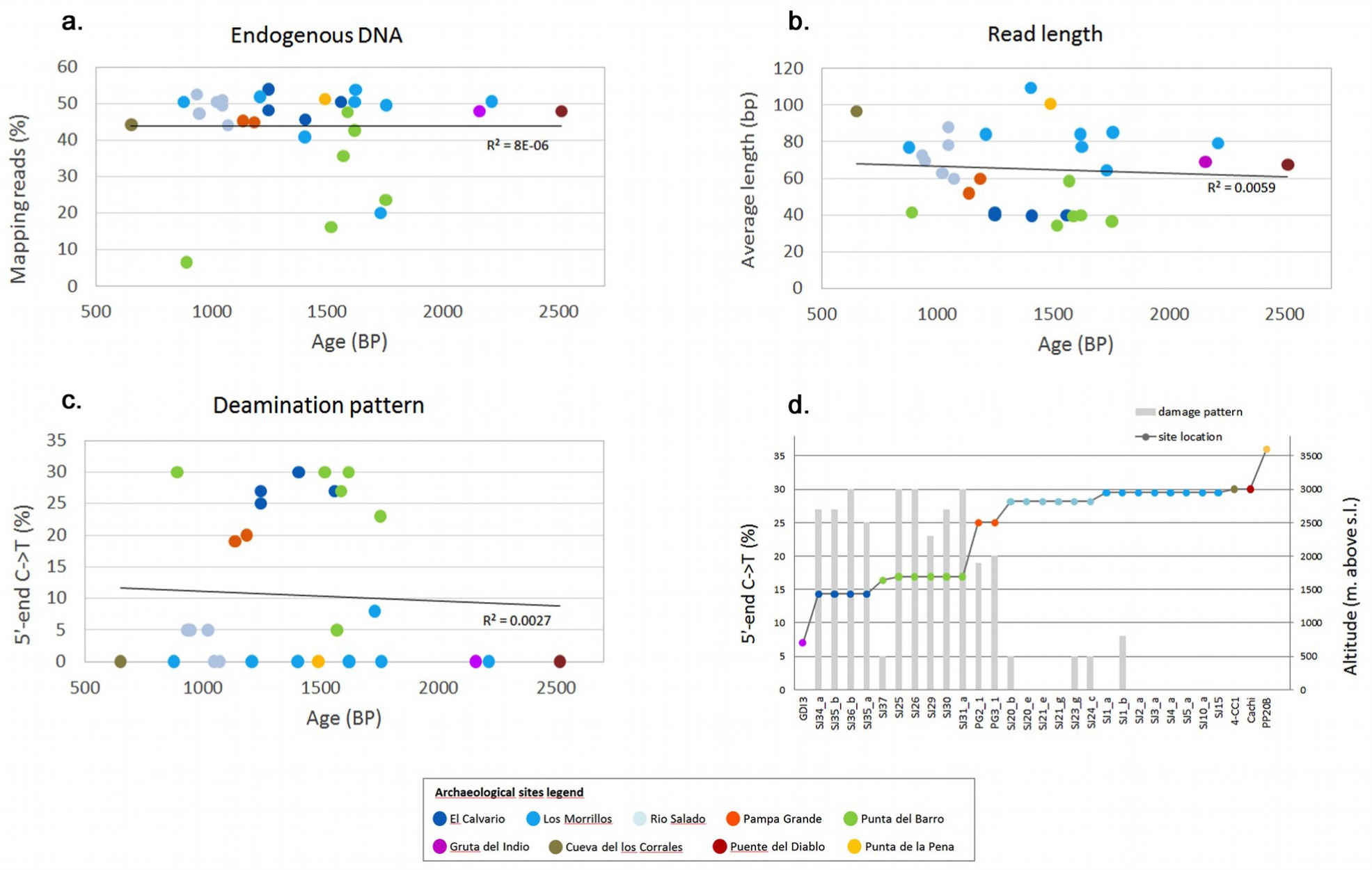
Endogenous DNA preservation in the 30 ancient bean seeds of the preliminary screening. a) Proportion of reads mapping to the common bean reference genome vs. age of the samples; b) Average read length vs. age of the samples; c) Misincorporation pattern at 5’ end vs. age of the samples; d) Misincorporation pattern at 5’ end vs. site locations.”

### S6. Modern accessions genomic data

Genomic data from modern domesticated and wild common bean samples were included in our analyses (Table S2). In particular, we included: *i*) 12 modern domestic common bean genomes from both Andean and Mesoamerican domestication gene pools, sequenced by the ERA-CAPS funded project “BeanAdapt: The Genomics of Adaptation during Crop Expansion in Bean” (unpublished; provided by the coordinator of the project, Prof. Roberto Papa, Marche Polytechnic University); *ii*) two published genomes from wild *P. vulgaris* (G19901 from Argentina and G24594 from Chiapas, Mexico; Rendòn-Anaya et al 2017); *iii*) one published genome of the wild relative *P. hintonii* to be used as outgroup in our selection scan (Rendòn-Anaya et al 2017). Trimmomatic v0.36 (Bolger et al 2014) was used to remove poor-quality regions from downloaded Illumina reads, discarding nucleotides having an average base quality score less than 20 across a 5bp window. Reads longer than 35bp were mapped to the *P. vulgaris* reference genome v2.1 in the paired-end mode using bwa mem (Li and Durbin 2009) with the parameter “-M”. The quality of bam alignments was improved removing PCR duplicates with Picard (software.broadinstitute.org) and performing the Indel realignment procedure, as described above for ancient samples. Depth of coverage was estimated in each sample using Samtools depth.

**Table S2.** Modern genomes included in our analyses. Gene pool indicates to which gene pool the sample belongs; AD: Andean domesticated, MD: Mesoamerican domesticated, AW: Andean wild, MW: Mesoamerican wild.

| Sample ID | Manuscript ID | Species | Wild/<br>Domestic | Gene<br>pool | Country/Sampling<br>site | Source |
| --- | --- | --- | --- | --- | --- | --- |
| ECa005 | CHI1 | *P. vulgaris* | D | AD | Chile | Bean Adapt Project |
| ECa006 | BOL1 | *P. vulgaris* | D | AD | Bolivia | Bean Adapt Project |
| ECa016 | CHI2 | *P. vulgaris* | D | AD | Chile | Bean Adapt Project |
| ECa038 | PER1 | *P. vulgaris* | D | AD | Perú | Bean Adapt Project |
| ECa040 | ARG1 | *P. vulgaris* | D | AD | Argentina | Bean Adapt Project |
| ECa041 | ARG2 | *P. vulgaris* | D | AD | Argentina | Bean Adapt Project |
| ECa086 | MEX1 | *P. vulgaris* | D | MD | México | Bean Adapt Project |
| ECa095 | MEX2 | *P. vulgaris* | D | MD | México | Bean Adapt Project |
| ECa104 | MEX3 | *P. vulgaris* | D | MD | México | Bean Adapt Project |
| ECa332 | ARG3 | *P. vulgaris* | D | AD | Argentina | Bean Adapt Project |
| ECa330 | ARG4 | *P. vulgaris* | D | AD | Argentina | Bean Adapt Project |
| ECa333 | BOL2 | *P. vulgaris* | D | AD | Bolivia | Bean Adapt Project |
| G24594 | MEX-W | *P. vulgaris* | W | MW | México | SRX540122 |
| G19901 | ARG-W | *P. vulgaris* | W | AW | Argentina | SRX2537721 |
| *P. hintonii* | - | *P. hintonii* | W | - | México | SRX2537735 |

### S7. Analysis of genomic diversity

To test whether there has been a decrease in genetic diversity due to the history of domestication, we compared levels of diversity between ancient and modern common bean seeds both as individual- and as population-level diversity. Genetic diversity was measured estimating Watterson’s Theta (*θ*_*w*_) in non-overlapping windows of 10kb, taking into account per-individual inbreeding coefficients and hence accounting for possible deviations from Hardy-Weinberg equilibrium, as proposed by Vieira et al 2013. This is a two-step procedure: first, ANGSD v0.916 (Korneliussen et al 2014) was used to call a high confidence SNP set based on GATK genotype likelihoods (-GL 2) including only alignment positions having an associated base and mapping quality score equal or higher than 20 and 30, respectively. To increase the reliability of the called set, we set the baq computation (-baq 1), adjusted the mapping quality in case of excessive mismatches (-C 50), and retained only positions having a high probability to be polymorphic (-SNP_pvalue 1e-6), without any missing data. From this set, we randomly sampled 2,000 SNPs without replacement and we used ANGSD to recompute the genotype likelihoods at these positions (-doGlf 3) applying the same quality filters. The program ngsF (Vieira et al 2013) was then used to estimate the individual genome-wide inbreeding coefficient for each accession using default parameters, except for the –min_epsilon option that was decreased to 10^−9^. The SNP sampling and the estimation of inbreeding processes were repeated five times in order to obtain five independent estimates of the inbreeding coefficient for each individual. The average across replicates was then used as point estimate of each individual inbreeding coefficient.

We then estimated a site frequency spectrum, using the command -doSaf 2 in ANGSD, to calculate the per-site posterior probabilities of the site allele frequencies based on individual genotype likelihoods accounting for the previously estimated inbreeding coefficients. Genotype likelihoods were calculated with the GATK method (– GL 2 option), and a folded spectrum was used (*i*.*e*., unknown ancestral state, option –fold 1). Options -minQ 20 -minMapQ 30 -setMaxDepth 50 were applied to filter low quality bases and positions showing excessive coverage, while the option -sites was set to filter for *callable* regions. The option -noTrans 1 was applied to all the above steps to systematically remove transitions from the analysis. The output of this command line was then used as input for the calculation of per-site thetas, using the command -doThetas 1. Window-based theta estimates were calculated using the command thetaStat do_stat. Windows having a proportion of usable sites less than 50% were discarded.

**Table S3.** Estimates of diversity in ancient and modern accession of common bean from Central (Meso) and South (Andean) America. The number of 10kb-windows with at least 50% of the region covered is reported.

| Manuscript ID | Gene pool | Country | 10kb-windows | $\theta_w$ mean | $\theta_w$ median |
| --- | --- | --- | --- | --- | --- |
| CHI1 | Andean | Chile | 24486 | 1.29E-05 | 9.62E-08 |
| BOL1 | Andean | Bolivia | 24686 | 1.56E-05 | 0 |
| CHI2 | Andean | Chile | 24493 | 1.18E-05 | 1.73E-07 |
| PER1 | Andean | Perú | 24597 | 9.62E-06 | 8.87E-08 |
| ARG1 | Andean | Argentina | 24636 | 1.51E-05 | 1.18E-08 |
| ARG2 | Andean | Argentina | 23827 | 7.44E-06 | 2.98E-07 |
| MEX1 | Meso | México | 21749 | 4.66E-05 | 9.05E-07 |
| MEX2 | Meso | México | 20071 | 2.42E-05 | 1.06E-06 |
| MEX3 | Meso | México | 20965 | 3.86E-05 | 5.76E-07 |
| ARG3 | Andean | Argentina | 23674 | 8.96E-06 | 1.40E-06 |
| ARG4 | Andean | Bolivia | 24272 | 1.67E-05 | 2.35E-06 |
| BOL2 | Andean | Argentina | 24491 | 1.99E-05 | 4.43E-07 |
| 15-CC | Andean | Argentina | 15865 | 3.02E-05 | 1.72E-05 |
| 09-CM | Andean | Argentina | 13284 | 3.78E-04 | 3.44E-04 |
| 01-PD | Andean | Argentina | 20897 | 5.48E-05 | 5.41E-07 |
| 03-GI | Andean | Argentina | 15241 | 1.79E-04 | 1.50E-04 |
| 07-PP | Andean | Argentina | 15303 | 3.40E-05 | 1.03E-05 |
| 14-CM | Andean | Argentina | 9372 | 1.12E-04 | 9.15E-05 |
| 04-CM | Andean | Argentina | 9892 | 8.31E-05 | 6.10E-05 |
| 10-CM | Andean | Argentina | 8422 | 1.80E-04 | 1.50E-04 |
| 08-CM | Andean | Argentina | 13130 | 6.91E-05 | 4.27E-05 |
| 12-CM | Andean | Argentina | 16859 | 1.15E-04 | 9.23E-05 |
| 11-CM | Andean | Argentina | 18346 | 5.08E-05 | 1.25E-05 |
| 13-CM | Andean | Argentina | 22600 | 8.76E-05 | 3.10E-05 |
| 02-CM | Andean | Argentina | 21224 | 4.99E-05 | 8.47E-06 |
| 06-PB | Andean | Argentina | 19426 | 5.51E-04 | 5.17E-04 |
| 05-CM | Andean | Argentina | 17519 | 1.85E-04 | 1.58E-04 |
| ARG-W | Andean wild | Argentina | 24255 | 4.08E-05 | 1.93E-06 |
| MEX-W | Meso wild | México | 22583 | 1.51E-03 | 9.25E-04 |

### S8. Re-calibrating estimates of modern genomic diversity

Genomic diversity of each modern accession was likely reduced by the protocols commonly used before whole-genome sequencing, which included two cycles of selfing reproduction after collecting the seeds from the seed bank. In addition, previous maintenance practices applied at each seed bank could have also affected genome-wide diversity. To test whether such very recent events could be the cause of the observed pattern of lower diversity in modern seeds as compared to ancient seeds, we re-estimated whole-genome diversity in pairs of seeds from closely related cultivars. The rationale behind this test is the following. Diversity loss caused by recent seed bank or sequencing protocols occurred at random along the genome in each cultivar, so that each cultivar should have lost a different fraction of its diversity. Taking together two (or more) closely related cultivars should restore most of the variation prior to the stocking into the seed banks. Of course, this estimate is also affected by the possible genetic divergence between the cultivars used in each pair.

Estimates of diversity (*θ*_*w*_) in pairs of cultivars should better represent the single cultivar variation before seed bank and sequencing protocols (or even over-estimate, due to cultivar genetic divergence). We selected the two modern Chilean cultivars, already included in all our analyses (see Table S2) and added three cultivars from the same country sequenced within the ERA-CAPS funded project “BeanAdapt: The Genomics of Adaptation during Crop Expansion in Bean” (unpublished; provided by the coordinator of the project, Prof. Roberto Papa, Marche Polytechnic University). We estimated *θ*_*w*_ in ten random pairs of seeds. Specifically, we compared average *θw* estimated in three groups of accessions: 10 pairs of Chilean accessions, 12 single modern accessions from South America, and in 15 ancient seeds. Alongside individual variation, we also compared the distribution of *θw* across all 10kb-windows from each group.

As expected, genome wide diversity in pairs of seeds is higher than in single seeds (Table S4). However, average modern seed pairs diversity is still more than three times lower (Mann-Whitney U test, *P* = 0.003) than that estimated in ancient seeds (Table S3). The same pattern appears when considering the summarized diversity across all 10kb-windows in each group (Figure S4).

**Table S4.** Estimates of diversity in five accessions from Chile (two of them were also used in the main analyses) and in the ten pairs of Chilean samples. The number of 10kb-windows with at least 50% of the region covered is reported.

| Manuscript ID | 10kb-windows | $\theta_w$ mean | $\theta_w$ median |
| --- | --- | --- | --- |
| CHI1 (ECa005) | 24486 | 1.29E-05 | 9.62E-08 |
| CHI2 (ECa016) | 24493 | 1.18E-05 | 1.73E-07 |
| CHI3 (ECa004) | 24524 | 1.37E-05 | 6.18E-08 |
| CHI4 (ECa018) | 24609 | 1.57E-05 | 3.32E-08 |
| CHI5 (ECa020) | 24541 | 1.47E-05 | 9.38E-08 |
| CHI3.CHI1 | 27870 | 4.33E-05 | 2.89E-07 |
| CHI3.CHI2 | 27875 | 4.24E-05 | 4.43E-07 |
| CHI3.CHI4 | 27912 | 3.96E-05 | 1.95E-07 |
| CHI3.CHI5 | 27891 | 4.63E-05 | 3.46E-07 |
| CHI1.CHI2 | 27856 | 4.47E-05 | 4.89E-07 |
| CHI1.CHI4 | 27903 | 5.32E-05 | 3.14E-07 |
| CHI1.CHI5 | 27889 | 4.32E-05 | 3.69E-07 |
| CHI2.CHI4 | 27916 | 4.57E-05 | 4.07E-07 |
| CHI2.CHI5 | 27888 | 3.59E-05 | 4.49E-07 |
| CHI4.CHI5 | 27930 | 5.61E-05 | 4.56E-07 |

**Figure S4.**
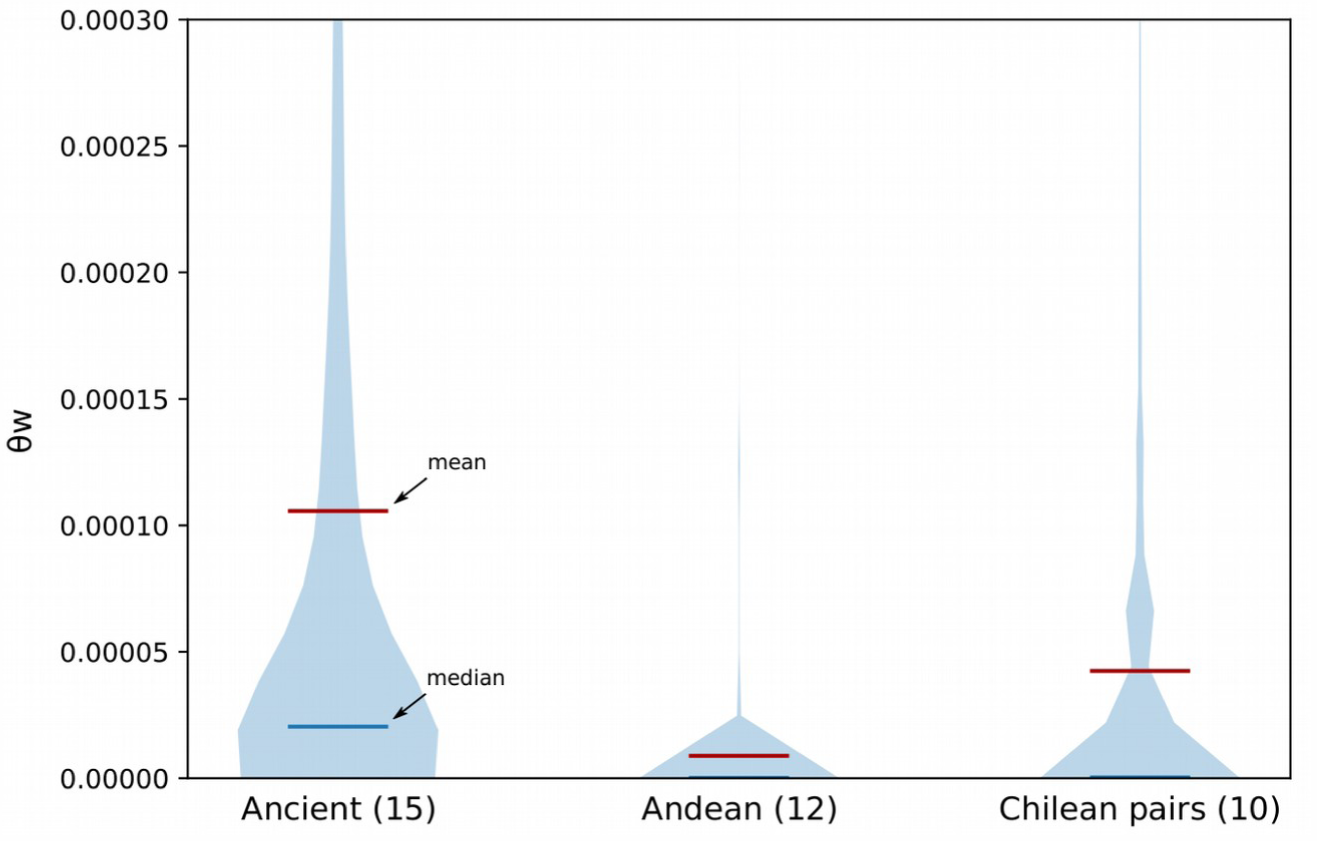
Estimates of whole-genome diversity in 15 ancient seeds (Ancient), in 12 modern accession from South America (Andean), and in 10 pairs of cultivars from the same region (Chile). Distribution of *θ*_*w*_ are based on all 10kb-windows across all samples in each group.

### S9. Structure of the genomic diversity in ancient beans

A haploid calling on all samples (15 ancient, 15 modern) was performed on each chromosome, separately, using ANGSD 0.916 (Korneliussen et al 2014). We set -minQ 20 and -minMapQ 30 to exclude low quality bases and low mapping quality bases; -only_proper_pairs 0 to use all reads (not only reads with both mapping pairs); -dohaplocall 2 to select the most frequent base in the haploid calling (*i*.*e*., not a random one); -doCounts 1 to count the bases at each sites after filters; -minMinor 2 to exclude singletons and -noTrans 1 to exclude transitions. The output of the haploid calling was then masked according to the three genomic regions as defined above (*callable, neutral* and *exons*) using *Bedtools intersect* (Quinlan and Hall 2010).

The main components of the genetic diversity and the eventual admixture proportion in all samples were inferred by *Admixture* analyses (Alexander and Novembre 2009). This analysis was performed using the haploid data after combining all chromosomes in one file. As *Admixture* input is binary *Plink* (.bed) or ordinary *Plink* (.ped), the haploid data was converted in tped format (*Plink* transposed text genotype table), using the haploToPlink function implemented in *ANGSD*. Non biallelic sites were excluded. This file was then converted to bed format, using –make-bed option in plink-1.90 (Chang et al 2015). The option –bp-space 50000 was added in order to select one SNP every 50kb (thinning) and minimize the effect of linkage disequilibrium in the *Admixture* analysis. We tested *k* values from 1 to 15, and cross validation error (Alexander and Lange 2011) was calculated for each *k* value, adding --cv flag to the command line. Cross validation plots and admixture plots (for each K value) were finally created in *R*.

**Figure S5.**
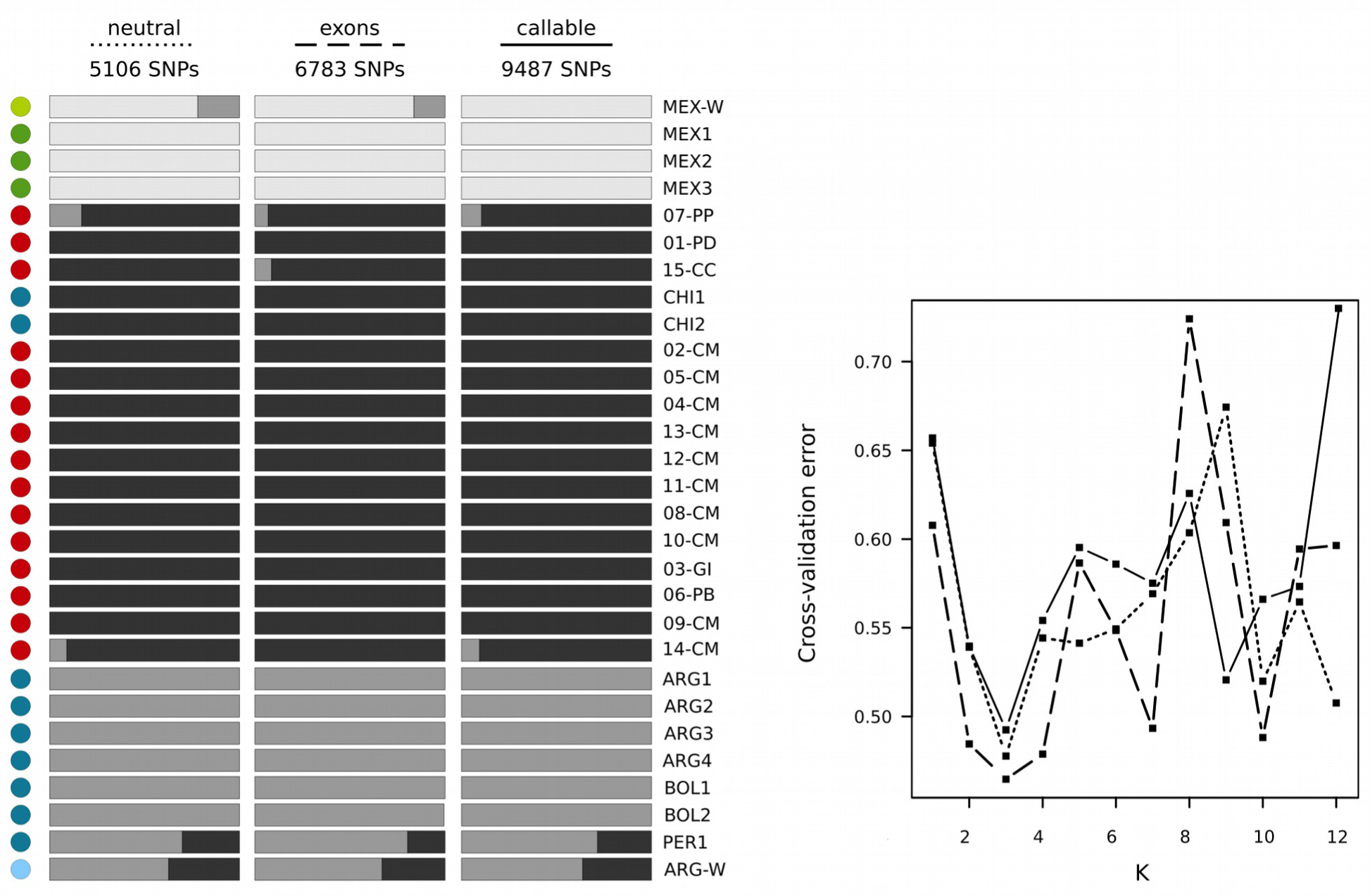
Structure of the genetic diversity inferred using three different genomic regions (*callable:* solid line, *exons:* large dashed line, *neutral:* small dashed line). For all genomic regions *k* = 3 represented the nest fitting model. Ancient domesticated Andean (red), modern domesticated Andean (blue) and Mesoamerican (green), wild Andean (pale blue) and Mesoamerican (pale green).

The output of the haploid calling masked according to the three genomic regions was also used to calculate per chromosome pairwise genetic distances across all modern and ancient individuals using a custom *awk* script (available upon request). Per-chromosome distances were then combined into a single genome-wide distance matrix, which was used to reconstruct a Neighbor-Joining tree using the *ape* package in *R* (Paradis et al 2004). Trees were visualized with FigTree (http://tree.bio.ed.ac.uk/software/figtree). As in the Admixture analysis, the three genomic regions produced almost identical results (the Neighbor-Joining tree based on callable region only is presented in Figure 1d in the main text).

Population structure was also inferred using the method implemented in FineStructure (Lawson et al 2012). The pattern of similarity and dissimilarity between haploidized individuals can be visualized by a distance matrix, and discrete populations can be separated by a cladogram. This method is able to capture at least as much information as that captured by other exploratory analyses such as PCA and Structure, but it also has the advantage to help identifying the pattern of population structure and to combine the information at linked markers. The latter advantage is of course reduced, or absent, in our haploidized data set. We used Plink to filter the dataset in order to exclude any variable position with missing data (not allowed in FineStructure) and the plink2chromopainter.pl conversion script to prepare the input file to run the FineStructure pipeline. In the first step of the pipeline, haplotype blocks are “painted” (Li and Stephens 2003) according to the nearest (in terms of genetic distance) individual in the sample (chromosome painting step). Painting results are then employed to build a distance matrix summarizing the blocks similarity between all pairs of individuals. The distance matrix is, in turn, the basis of a Bayesian clustering approach aiming at partitioning the sample into *K* homogeneous groups.

**Figure S6.**
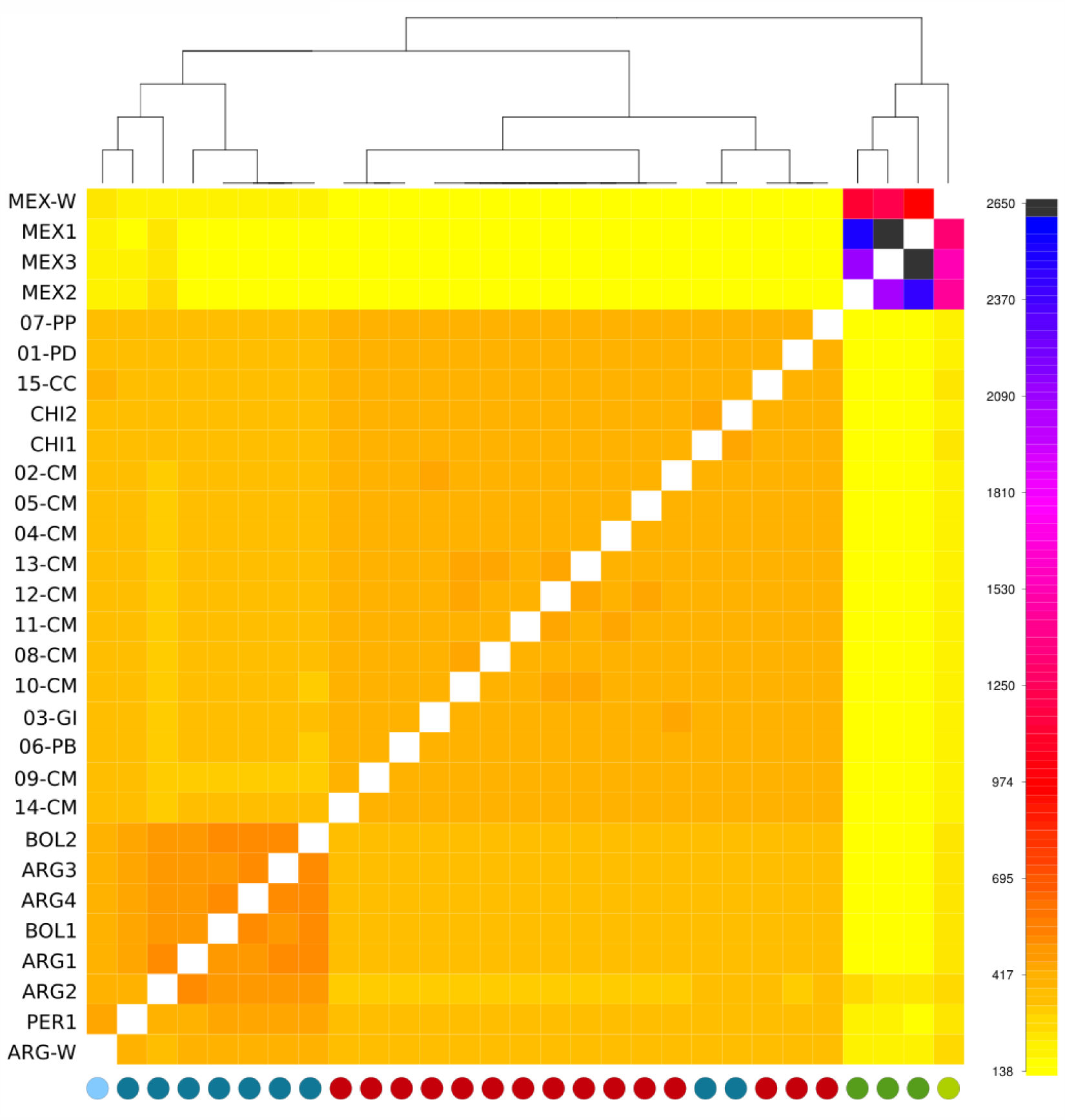
Finestructure analysis based on callable genomic region (483885 SNPs). Ancient domesticated Andean (red), modern domesticated Andean (blue) and Mesoamerican (green), wild Andean (pale blue) and Mesoamerican (pale green).

### S10. Genes-focused scan for signature of ancient and/or recent selection

A gene-by-gene selection scan was performed to identify genes targeted by artificial selection during early and/or late stages of domestication. A gene underlying a desirable trait is expected to have undergone one or more selective sweeps targeting specific alleles coupled with recurrent hitchhiking at linked neutral sites. Selected genes will then accumulate variants which are either directly advantageous or got fixed because of locally low effective population size due to linked selection (Slotte 2014, Renaut and Rieseberg 2015, Beissinger et al 2016). We then expect to find an excess of fixed differences at genes under selection when samples of pre (*i*.*e*., wild) and post domestication populations are compared.

Leveraging our ancient data, we tested each of the 27,000 genes for a significant enrichment of fixed alternative alleles in *i*) the ancient + modern Chilean samples as compared to wild samples (one wild Andean *P. vulgaris* and one *P. hintonii* as wild outgroup), representing the signature of ancient selection (before 2,500 years ago - topology AS; Fig. 2a in the main text) and in *ii*) the modern Chilean samples alone as compared to the ancient + wild samples, representing the signature of recent selection (after 600 years ago - topology RS; Fig. 2a in the main text).

To increase the number of linked neutral sites, we initially counted the fixed alternative alleles in genomic regions including a gene and 10 kb before and after it (*expanded genes*). A significant enrichment in each gene of SNPs showing either AS or RS topology was tested as follows. First, we estimated an expected distribution of fixed alternative alleles in any topology if SNPs were distributed at random across all *expanded genes*. In particular, after shuffling all SNPs found across all *expanded genes*, we assigned them equally to 27,000 groups, corresponding to pseudo *expanded genes*, and estimated the distribution of SNPs in each possible topology. Such distributions were then used to test the significant enrichment of SNPs in one of the possible topologies in each real *expanded gene*. We calculated the exact *p*-value of a disproportional accumulation of SNPs in any of the tested topologies for each *expanded gene* by means of a binomial test: *p*-values were then transformed into *q*-values (False Discovery Rate, FDR) using the *python* function qvalue.py (https://github.com/nfusi/qvalue/blob/master/qvalue/qvalue.py) based on (Storey and Tibshirani, 2003); only *expanded genes* with a FDR<0.001 were considered as significantly enriched in a particular topology.

The final lists of *expanded genes* enriched for either AS or RS (or both) were further sorted by the number SNPs with fixed alternative alleles falling inside or within 1kb from the gene. Applying a further stringent criterion, genes (± 1kb) showing less SNPs with fixed alternative alleles than the 99-percentile of all 27,000 genes were discarded. The two final lists of genes showing signature of AS and/or RS were grouped into functional clusters using the web-based software David (Huang et al 2009). For this analysis, we could only use genes with a known ortholog in *A. thaliana*. We used the functional enrichment score estimated by David only to rank the functional clusters found in the two lists of genes without excluding any cluster. Analyses were replicated using the genome of a wild Mesoamerican *P. vulgaris* as outgroup instead of *P. hintonii* (Table S5).

